# Network rewiring of a pneumococcal communication system promotes stress adaptation in a globally successful lineage

**DOI:** 10.64898/2025.12.10.693554

**Authors:** Bárbara Ferreira, Carina Valente, Ozcan Gazioglu, Hasan Yesilkaya, N. Luisa Hiller, Raquel Sá-Leão

## Abstract

Cell-cell communication (CCC) systems regulate bacterial behaviors and shape social interactions within microbial communities. In *Streptococcus pneumoniae*, several CCC systems coordinate intra- and inter-strain signaling, yet the genetic plasticity of their components and regulons remains underexplored. Here, we investigate a divergent allelic variant of the TprA/PhrA CCC system, a regulator-pheromone pair widely conserved in *S. pneumoniae*. We characterize PhrA1.2, the second most prevalent PhrA pheromone, as a non-signaling variant found in the globally distributed GPSC6 lineage, with presence in isolates dating back to the 1960s. In GPSC6, PhrA1.2 co-occurs with TprA1.2, a truncated transcriptional regulator carrying premature stop codons that eliminate the predicted pheromone-binding domain. Functional assays show that PhrA1.2 is non-functional, both when paired with the truncated TprA1.2 and when paired with the canonical regulator allele TprA1.1. Unexpectedly, despite its truncation, TprA1.2 activates genes involved in oxidative stress responses, including those related to iron-sulfur cluster formation and redox homeostasis. Loss of *tprA1.2* impairs growth and increases hydrogen peroxide sensitivity, while *in vivo* experiments reveal a fitness defect during nasopharyngeal colonization. Together, our findings show that the TprA1.2 allele represents a regulator that has lost its communication component while modifying its regulon. This variant promotes adaptation to oxidative stress and contributes to colonization, demonstrating circuit rewiring of a peptide-regulator CCC system in an epidemiologically relevant *S. pneumoniae* lineage.

## INTRODUCTION

*Streptococcus pneumoniae*, or the pneumococcus, asymptomatically colonizes the human upper respiratory tract, particularly in young children (1). Despite its role as a frequent commensal, it remains a major human pathogen responsible for a wide variety of infections such as otitis media, pneumonia or meningitis (2, 3). The widespread use of pneumococcal conjugate vaccines (PCVs) has substantially reduced the burden of vaccine-type disease (4, 5), but has also driven major shifts in pneumococcal population structure, favoring the emergence of non-vaccine serotypes and the persistence of multidrug-resistant lineages (3, 6, 7).

The global pneumococcal sequence cluster 6 (GPSC6), historically known as PMEN3 clone or Spain9V-ST156, is one of the most successful and persistent clones in the post-PCV era (8, 9). First identified in Spain in the late 1980s, GPSC6 was initially linked to serotype 9V, and rapidly became a major cause of invasive pneumococcal disease through β-lactam non-susceptibility mediated by mosaic *pbp* genes (10). A capsular switch to serotype 14 in the 1990s allowed global spread (9, 10). Subsequent diversification through recombination produced variants expressing non-vaccine serotypes such as 11A, 15B/C, 19A, 23A, 24F, and 35B (7, 11–14). Worldwide long-term surveillance studies have consistently demonstrated the remarkable persistence and adaptability of GPSC6 across multiple vaccine eras. In Spain, GPSC6 remained a major cause of invasive pneumococcal disease (IPD) for over three decades (1987-2016), maintaining circulation despite vaccine-driven serotype replacement and the decline of other resistant clones (7, 15). In Brazil, GPSC6 was the main pneumococcal lineage in children IPD data from pre and pos-PCV10 implementation (16). In Portugal, carriage studies showed that GPSC6 was already established before PCV implementation (17) and remained one the dominant multidrug-resistant lineages throughout the PCV7 and PCV13 periods (11, 18).

Genomic analysis revealed that GPSC6 has a high transformation potential and has undergone recurrent recombination at key loci such as *pbp2x*, *pbp1a*, and *murM*, as well as frequent acquisition of macrolide and tetracycline resistance elements (7, 19, 20). Many GPSC6 isolates also carry the pilus islet-1, which enhances adhesion and persistence in the nasopharynx (21, 22). Together, these findings establish GPSC6 as a paradigm of pneumococcal resilience and adaptability, capable of maintaining circulation in both colonization and invasive disease despite sustained vaccine and antibiotic pressures.

Bacterial cell–cell communication (CCC) systems are critical for coordinating population-level behaviors such as competence, antimicrobial production, and virulence (23). In Gram-positive bacteria, CCC often relies on secreted peptide signals sensed by membrane-bound or cytoplasmic receptors that regulate gene expression (23). Among the latter, the RNPPA is the main superfamily, it includes peptide-mediated regulation via cytoplasmic transcription factors and is common across streptococci (24, 25). The pneumococcus includes multiple members of the RRNPPA, including alleles of the Tpr/Phr systems (26).

In *S. pneumoniae*, three *tpr/phr* cassettes have been identified, *tprA/phrA, tprB/phrB*, and *tprC/phrC*. These are homologous to the well characterized PlcR/PapR system from *Bacillus* (27, 28). The best-characterized cassette, *tprA/phrA*, regulates a downstream putative lantibiotic locus and influences colonization and disease (28–30). In the *S. pneumoniae* strain D39, TprA functions as a transcriptional factor and represses the downstream genomic region, which includes its cognate *phrA* pheromone gene. The PhrA peptide is secreted as a precursor, processed during export via the Sec secretion pathway, and re-imported into the cytoplasm through the Ami oligopeptide transporter, at high cell-density. Once internalized, PhrA binds to cytoplasmic TprA, causing its release from the promoter. This leads to the de-repression of *phrA* and the lantibiotic genes, resulting in strong activation of their expression (28).

While PhrA-mediated activation has been established in a reference strain, less is known about how this CCC systems varies across natural pneumococcal populations. In particular, the contribution of lineage-specific genetic variation - whether in peptide, receptor, or promoter sequences - to the functional diversification or silencing of CCC remains poorly understood. Previous work from our group identified a lineage-specific TprA2/PhrA2 cassette in the PMEN1 lineage, marked by unidirectional signaling with the TprA paralogue. The co-variance between receptor and peptide suggests co-evolution between Tpr receptors and Phr pheromones (31). These findings raise the question of whether other cassettes across diverse pneumococcal backgrounds display altered or even non-canonical behavior.

Here, we report the genomic and functional characterization of a non-canonical *tprA/phrA* pherotype in *S. pneumoniae*, defined by a variable allele of PhrA, PhrA1.2, and a truncated version of the TprA1.2 regulator. Predominantly found in the globally disseminated GPSC6 lineage, this system has lost the cell-cell communication capacity yet retains regulatory activity over genes involved in oxidative stress. Functional assays, transcriptomics, and *in vivo* experiments revealed that TprA1.2 contributes to bacterial fitness and colonization through pheromone-independent regulation. Our findings highlight how a conserved CCC module has been evolutionarily reconfigured - via domain truncation and promoter variation - to support lineage-specific adaptation in a major pneumococcal clone.

## MATERIAL AND METHODS

### Bacterial strains, peptides and growth conditions

The bacterial strains and peptides used in this study are listed in **Table S1**. For routine growth, strains were streaked from skim milk-tryptone-glucose-glycerine, STGG, glycerol stocks onto Tryptic Soy Agar with 5% sheep blood (BAP, BBL Stacker Plates, BD) and incubated overnight at 37°C in 5% CO_2_. For all assays, pre-inocula were prepared by growing bacteria in C+Y_YB_ medium (for transformation protocols, (32)) or in chemically defined medium supplemented with galactose (CDMgal, (33)) for all other protocols. Cultures were grown to an optical density at 600 nm (OD_600_) of 0.5 and stored at -80°C in 15% glycerol.

### Comparative genomics and *in silico* functional prediction of PhrA1.2/TprA1.2 pherotype

To characterize the genomic context of *phrA1.2* in *S. pneumoniae*, we analyzed a curated dataset of 7,548 *S. pneumoniae* genomes (34) that we refer to as the GoldenSet. For each strain positive for *phrA1.2*, we examined the corresponding *tprA* allelic variant, the downstream gene region (previously shown to play a role in the regulation of gene expression by the *tprA/phrA* system), and the epidemiological background of the strain. Eacsh of these strains was annotated with its Global Pneumococcal Sequence Cluster (GPSC) using PathogenWatch (http://pathogen.watch, (35, 36)) to explore potential epidemiological associations. The same approach was done for a colonization collection from 2018-2020 isolates from Portugal (11).

Comparative sequence analyses of TprA variants were performed using Qiagen CLC Genomics Workbench software version 9.5.1 (Qiagen), and protein domain prediction for TprA was conducted using NCBI Conserved Domain Search (37), InterProScan tool’s (38). Structural models of selected TprA variants were generated using AlphaFold v3.0 (39) to further infer potential functional differences. All DNA and protein alignments were performed under default parameters unless otherwise stated.

For promoter analysis of *phrA*, P*phrA*, we extracted the region downstream of *tprA* and categorized the promoter sequences based on single-nucleotide differences relative to the reference promoter from the model *S. pneumoniae* strain D39, *PphrA1.1*.

### Construction of markerless genetically modified strains

Deletion of *tprA* and restoration of its full-length coding sequence were performed via site-directed homologous recombination using a two-step transformation strategy incorporating the Fluorescent Removable Antibiotic Cassette (FRANC) system (40). In the first transformation step, the FRANC cassette - containing the spectinomycin resistance gene (*aad9*), the sucrose sensitivity gene (*sacB*), and the red fluorescent marker *mCardinal* - was inserted at the target genomic locus, either to disrupt *tprA* (for deletion) or to replace a modified region (for site-directed mutagenesis). Colonies with successful cassette insertion were selected based on spectinomycin resistance, red fluorescence (mCardinal), and PCR verification. In the second transformation step, the cassette was replaced via homologous recombination using donor DNA containing either (i) the upstream and downstream flanking regions (for clean deletion) or (ii) flanking regions plus the desired modified sequence (for domain restoration or amino acid substitution). For restoration of the full *tprA* coding sequence, donor DNA was synthesized using the wild-type *tprA* sequence from the D39 strain. A list of primers used for the construction of markerless genetically modified strains is provided in **Table S1.**

For transformation, bacterial pre-cultures were grown in C+Y_YB_ medium (pH 7.4) at 37 °C until reaching an OD_600_ of 0.5. Cultures were then diluted 1:100 into fresh C+Y_YB_ medium and regrown to an OD_600_ of 0.1. At this point, 150 ng of the donor DNA (amplicon) and 125 ng of synthetic competence-stimulating peptide 1 (GeneScript, USA; **Table S1**) were added to each culture. The mixtures were incubated at 37 °C for 4 hours to allow transformation. After incubation, cultures were plated onto blood agar plates supplemented with 100 µg/mL spectinomycin for selection. Plates were incubated overnight at 37 °C in 5% CO_2_. Resulting colonies were screened by colony PCR to confirm correct genomic integration of the desired construct.

### DNA extraction for whole genome sequencing

Pure cultures of GPSC6 TprA1.2-Full and GPSC6 Δ*tprA1.2* were streaked onto tryptic soy agar (TSA) plates supplemented with catalase (∼240 U/mL) and incubated overnight at 37 °C in 5% CO₂. Bacterial lawn was collected by adding 500 μL of sterile phosphate-buffered saline (PBS). Of this culture, 200 μL was transferred to a tube containing 200 μL of lysis buffer (MagNA Pure Compact Nucleic Acid Isolation Kit, Roche Diagnostics GmbH) and 17.4 μL of RNase A (final concentration: 100 mg/mL). Samples were incubated at 37°C for 20 minutes. Genomic DNA was then extracted using the MagNA Pure Compact Nucleic Acid Isolation Kit (Roche Diagnostics GmbH) according to the manufacturer’s instructions.

DNA quality and purity were assessed by measuring A_260_/A_280_ and A_260_/A_230_ ratios using a NanoDrop spectrophotometer (ThermoFisher Scientific), along with visual evaluation of integrity by agarose gel electrophoresis. DNA concentration was determined using the Qubit dsDNA High Sensitivity Assay Kit (ThermoFisher Scientific), following the manufacturer’s protocol.

### Whole genome sequencing

Whole genome sequencing was done at the Genomics Unit of the Instituto Gulbenkian de Ciências (Oeiras, Portugal). Libraries were prepared using the Nextera XT DNA Library Preparation Kit (Pico input) and sequenced on an Illumina NextSeq platform, targeting a mean coverage depth of 100X. Raw paired-end reads were assembled and subjected to quality control using INNUca v4.2.2 (https://github.com/B-UMMI/INNUca), with default parameters.

Assembly pipelines were adjusted to account for the bacterial species and expected genome size, which were used as filtering criteria during quality control and contig inclusion/exclusion. Assembled genomes were annotated using Prokka pipeline version 1.13.3 (41).

### RNA extraction and qRT-PCR

Pre-cultures were grown in CDMgal medium at 37 °C with 5% CO₂ until they reached an OD_600_ of approximately 0.1, corresponding to early exponential phase. Cultures were then either split into two experimental conditions: one treated with 1 mM synthetic PhrA peptide (either PhrA1.1 or PhrA1.2), or DMSO as control (**Table S1**). After a 2-hour incubation at 37°C, cultures were immediately put on ice, and total RNA was extracted following a previously described protocol (42).

To ensure the absence of genomic DNA contamination, PCR was performed using *gapdh*-targeted primers with RNA as the template. For qRT-PCR, complementary DNA (cDNA) was synthesized from RNA using the Bioline cDNA Synthesis Kit, and amplification was carried out with the SensiFAST SYBR Kit (Bioline) on a CFX96 Touch Real-Time PCR System (Bio-Rad).

Experiments were performed with three independent biological replicates, each run in technical duplicate. Primer sequences are listed in **Table S1**. Data analysis was conducted using Bio-Rad CFX Manager software. Relative gene expression was normalized to the housekeeping gene *16S*, and fold changes were calculated using the 2^−ΔΔCT method (43).

### Transcriptional reporter assays

Bacterial strains were initially cultured overnight on blood agar plates (BAP) and then inoculated into fresh CDMgal medium. Once cultures reached an OD_595_ between 0.05 and 0.1, they were split into two conditions: one treated with DMSO (control) and the other with 1 μM of synthetic PhrA1.1 or PhrA1.2 peptide (**Table S1**). Growth and luminescence were monitored over a 6-hour period. At each hour, 150 μL of culture was transferred to white 96-well plates (Greiner, USA) for luminescence measurement using a Tecan Infinite M Plex plate reader (1000 ms integration time). In parallel, 700 μL of each culture was collected to measure optical density, serving as a control for growth differences across conditions.

### Bacterial growth assays

To compare fitness growth of wt and mutant strains, bacterial pre-cultures were grown in CDMgal medium to an OD_600_ of 0.5, then diluted 1:100 into fresh CDMgal supplemented with catalase (∼1600 U/mL). Cultures were incubated at 37 °C for 24 hours in 24-well polystyrene plates using an automated incubating plate reader (Tecan Infinite 200 Pro). Optical density at 595 nm was measured every 30 minutes. Prior to each measurement, orbital shaking with a 6 mm amplitude was applied to ensure adequate mixing and bacterial suspension. All experiments were performed independently on three separate days.

### Transcriptome anaylsis

Total RNA was extracted from three independent biological replicates of strains GPSC6wt, and GPSC6Δ*tprA1.2*. RNA-seq was performed by Novogene using the Illumina NovaSeq X Plus platform, generating paired-end reads with a minimum length of 150 bp and a depth of at least 20 million reads per sample. A strand-specific (directional) prokaryotic mRNA library was prepared for each sample, including ribosomal RNA depletion before sequencing. RNA sequencing data were analyzed following the workflow described by Pobre and Arraiano (44), with adaptations. Briefly, sequencing reads were aligned to the GPSC6wt reference genome using Bowtie2 (45), and alignment files were converted from SAM to BAM format using Samtools (46). Gene expression quantification and differential expression analysis were performed using the *edgeR* package in R. Genes were considered differentially expressed if they met the following criteria: false discovery rate (FDR) < 0.05, absolute log_2_ fold change > 1, and P-value < 0.001. To improve data reliability, additional manual curation was performed by excluding genes with fewer than 100 read counts. Functional annotation of selected genes was carried out using BLASTp searches of the corresponding protein sequences against the NCBI non-redundant protein database.

Genetic association networks of differently expressed genes were investigated using STRING v12. Since GPSC6 strain is not represented in STRING, each differentially expressed GPSC6 genes were first mapped to their closest orthologs in the reference *Streptococcus pneumoniae* R6 genome using BLASTp. The set of mapped orthologs was then submitted to STRING with the species restricted to *S. pneumoniae* R6. Default STRING parameters were used. Predicted functional partners returned by STRING that belonged to the same modules were retained to provide functional context. All networks were interpreted as orthology-based functional projections.

The transcriptomic data from this study were submitted to the European Nucleotide Archive (ENA) under accession number PRJEB103953.

### Hydrogen peroxide susceptibility assay

The susceptibility of GPSC6 strains to oxidative stress was assessed using hydrogen peroxide exposure, following the protocol described by Pericone *et al*. (47), with minor adaptations. Briefly, bacterial pre-cultures were grown in CDMgal medium at 37 °C until reaching an OD_600_ of approximately 0.3. Cultures were then divided into 100 μL aliquots. To each aliquot, an equal volume of either phosphate-buffered saline (PBS; control) or hydrogen peroxide was added to achieve final H_2_O_2_ concentrations of 20 mM or 40 mM in the treated samples.

The mixtures were incubated at 37°C with for 30 minutes. Following incubation, samples were serially diluted and plated onto blood agar plates. After overnight incubation, colony-forming units (CFUs) were counted to quantify viable bacteria. The percentage of surviving bacteria was calculated by comparing the CFU counts of H_2_O_2_ treated samples to those of the PBS-treated control.

### *In vivo* colonization model

The role of *tprA1.2* in promoting fitness during pneumococcal colonization was assessed using an established murine model (48). Female outbred mice (9–11 weeks old), bred and maintained at the Division of Biomedical Services, University of Leicester, were used for all experiments. Each mouse was intranasally inoculated with 5x10⁵ CFU of the bacterial strain of interest in 20 μL of phosphate-buffered saline (PBS, pH 7.0). Mice were anesthetized using 2.5% (v/v) isoflurane (Isocare) in oxygen delivered at a rate of 1.4–1.6 L/min. To minimize the risk of bacterial aspiration into the lungs, mice were maintained in a horizontal position during inoculation. The actual inoculum dose was confirmed by plating serial dilutions of the inoculum on blood agar plates. To quantify colonization, nasopharyngeal washes were collected from mice at 3 and 7 days post-inoculation. Bacterial loads were determined by plating serial dilutions of the washes on agar plates containing 5 μg/mL gentamicin.

## RESULTS

### A novel PhrA allele is highly enriched in the prevalent GPSC6 lineage

In a recent survey of the TprA/PhrA cell-cell communication (CCC) systems in *S. pneumoniae* we identified allelic variants of the PhrA pheromone (Ferreira et al., submitted). This analysis was performed using our GoldenSet, a database of 7,548 pneumococcal genomes (34) selected to capture the genomic diversity across the pneumococcal species.

We determined that one allele of PhrA, PhrA1.1, was dominant being encoded in >64.0% of the analysed strains. The next most common allele, PhrA1.2, was present in only 2.3% of strains (172 genomes of 7,548). The PhrA1.2 differs from the dominant PhrA1.1 by a single amino acid substitution in the active C-terminal region of the peptide (PhrA1.1: SNGLDV**V**KAD and PhrA1.2: SNGLDV**G**KAD). Although rare overall, PhrA1.2 was strikingly enriched in GPSC6, a lineage that has been globally disseminated since the 1990s and persisted despite widespread vaccine use targeting its dominant serotypes (7–9). Within the GoldenSet, 92.2% of GPSC6 genomes carried PhrA1.2 (165 of 179) (**Fig. 1A**). A few remaining instances were limited to 2.9% of GPSC39 genomes (3 of 102) and to all sampled genomes of GPSC268 (4 of 4, though this lineage is sparsely represented) (**Fig. 1A**, **Table S2**). Overall, 95.9% of all genomes encoding PhrA1.2 belonged to GPSC6 and almost all GPSC6 genomes encode PhrA1.2 highlighting its importance in this lineage.

**Figure 1.**
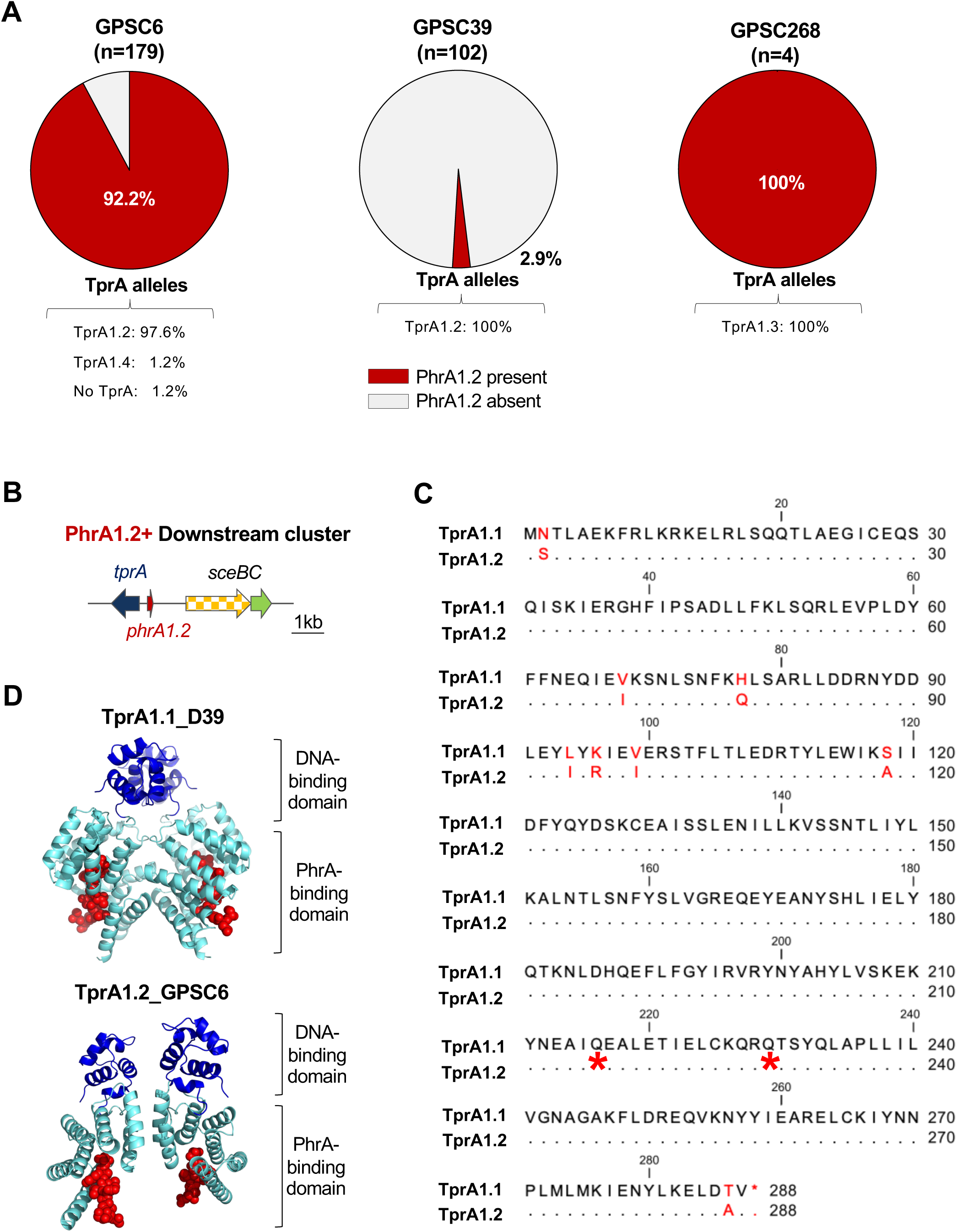
Distribution and characterization of the PhrA1.2 allele in *S. pneumoniae*. **A.** The prevalence of the PhrA1.2 allele was assessed across 7,548 curated pneumococcal genomes and detected in only three Global Pneumococcal Sequence Clusters (GPSCs): GPSC6, GPSC39, and GPSC268, with a strong enrichment in GPSC6. The corresponding TprA allelic variants present in strains carrying PhrA1.2 are also shown. Full amino acid sequences of these variants are provided in **Table S2**. **B.** Genomic organization downstream of *phrA1.2*. All strains harbour an incomplete *sce* bacteriocin locus. Genes are color-coded: *tprA* (blue, transcriptional regulator), *phrA* (red, pheromone), *sceB* (yellow, immunity protein), and *sceC* (green, ABC transporter). **C.** Comparison of TprA alleles. Alignment of TprA1.1 from strain D39 with TprA1.2, the predominant allele in GPSC6 PhrA1.2^+^ isolates (98.2%). Amino acid substitutions between the two proteins are shown in red. Premature stop codons at positions 216 and 229 are marked with red asterisks. **D.** Structural predictions of TprA1.1 and TprA1.2, in dimer conformations, in the presence of PhrA1.1 C10 or PhrA1.2 C10, respectively, as predicted by AlphaFold3. The TprA1.2 model shows truncation of the C-terminal PhrA-binding domain, while the DNA-binding domain remains intact.

Considering the presence of PhrA1.2 across GPSC lineages 6, 39, and 268, we examined with higher resolution the genetic relatedness of the clones within these lineages in the GoldenSet (**Table S3**). Our data shows that in GPSC6, PhrA1.2 has been present continuously since the early 1960s across multiple sequence types (STs) (**Table S3**). Despite this diversity, all PhrA1.2 positive isolates are associated with ST156, representing single-, double-, or triple-locus of this ST (**Table S4**). In contrast, in GPSC39, PhrA1.2 was only detected in genomes from the 1950s, with no representation in genomes from the later decades (**Table S3**); all 3 isolates belonged to the same ST. The limited number of isolates from GPSC268 precludes any conclusion on the prevalence or evolution of PhrA1.2 within this lineage.

To examine the distribution of this allele in recent years, we analyzed a dataset of 837 *S. pneumoniae* isolates collected from children attending Portuguese day-care centers between 2018 and 2020 (11). In this collection, GPSC6 accounts for 12.3% of isolates (107 of 873), exposing its current prevalence in the population. In accordance with the high frequency of PhrA1.2 in GPSC6, PhrA1.2 is present in 12.3% of the total dataset (100% of the GPSC6 isolates, **Table S5**). The elevated representation of GPSC6 in this period is consistent with parallel epidemiological surveys, which document the expansion of this lineage in Europe (7, 49, 50). Further, similar to the GoldenSet analysis, we also examined the genetic relatedness of clones within GPSC6 (the only lineage, in this setting, with PhrA1.2). As before, all STs identified within GPSC6 are related to ST156, representing single-, double-, or triple-locus of this ST (**Table S6**).

This close correspondence underscores a long-standing association between PhrA1.2 and ST156, a major clonal complex within GPSC6. Since its emergence in the 1990s as the Spain9V-ST156 clone (8, 10), this lineage has remained one of the most successful pneumococcal lineages through successive capsular switch events. Notably, the PhrA1.2 signaling cassette has been consistently present in ST156 genomes from the earliest available isolates, dating back to the 1960s, through to contemporary isolates from the 2020s (**Tables S2-S5**). Collectively, these findings establish PhrA1.2 as a defining and conserved feature of GPSC6.

### The PhrA1.2 allele co-occurs with a truncated version of the TprA regulator and an incomplete bacteriocin locus

PhrA peptides signal by binding their cognate regulator, TprA. To assess allele specificity across peptide-receptor pairs, we examined the distribution of TprA alleles co-occurring with PhrA1.2. In the GoldenSet, >95% of strains carrying PhrA1.2 (164 of 172) encoded a specific TprA variant, TprA1.2. The remaining genomes encoded TprA1.85 (4 isolates, 2.3%), TprA1.124 (2 isolates, 1.2%), or lacked TprA altogether (2 isolates, 1.2%) (**Fig 1A**). Within GPSC6, TprA1.2 was present in >97% of strains (161 of 165) (**Table S2**). Likewise, in the Portuguese dataset, all isolates with PhrA1.2 also encoded TprA1.2 (**Table S5**). Together, these findings demonstrate a consistent co-occurrence between PhrA1.2 and the TprA1.2 allele.

The TprA/PhrA system regulates a broad set of genes, but those with the strongest induction are located immediately downstream of the signaling locus (29, 40). In a recent survey, we found that the downstream region can encode diverse operons (Ferreira et al, submitted). Among them, two operons have been experimentally confirmed as TprA/PhrA-regulated bacteriocins: the *streptococcin E* (*sce*) locus (Ferreira et al, submitted) and the lantibiotic (*lan*) locus (29). We therefore examined the downstream region in strains encoding PhrA1.2. In both the GoldenSet and the Portuguese dataset, this region consistently corresponded to an incomplete *sce* operon, encoding the immunity protein, and ABC transporter, but lacking the genes for the active bacteriocin. (**Fig 1B**, **Table S2-3**).

We next compared the most prevalent TprA allele associated with PhrA1.2, TprA1.2, with TprA1.1, which has been validated experimentally (28). The alignment reveals two premature stop codons in TprA1.2 (**Fig 1C**), which lead to a partial loss of the tetratricopeptide repeats predicted to mediate PhrA binding (**Fig 1C**). The prediction of the PhrA binding domain is based on AlphaFold predictions and analogy with the crystal structure of the homologous systems, PlcR/PapR, in *Bacillus* (51). Thus, we deduce that the TprA1.2 allele is likely impaired in binding PhrA, raising the possibility that either the signaling system or the regulator is inactive in the GPSC6 lineage.

To evaluate structural differences between TprA alleles, we predicted the dimeric structures of TprA1.1 from D39 and TprA1.2 from GPSC6 in complex with their cognate PhrA peptide (**Fig. 1D**). For TprA1.1, a high-confidence model of the dimer was obtained, with PhrA1.1 consistently positioned at the C-terminal site for each monomer (pTM = 0.76, ipTM = 0.73). For TprA1.2, the resulting model predicted peptide binding, but the interface was less extensive and had lower confidence, reflected in reduced scores (pTM = 0.66, ipTM = 0.63). These results suggest that while both alleles may associate with their peptides, the interaction is predicted to be weaker or less stable in TprA1.2 compared with TprA1.1.

In summary, the GPSC6 lineage is marked by a distinctive signaling module: the PhrA1.2 pheromone and its co-occurring, truncated TprA1.2 receptor, accompanied by an incomplete *sce* operon. This organization suggests functional changes in the regulatory circuit of TprA/PhrA in GPSC6.

### The PhrA1.2 peptide does not regulate TprA variants

To assess PhrA1.2-dependent signaling in its native genetic context, we quantified gene expression responses to PhrA1.2 and PhrA1.1 using qRT-PCR in the GPSC6 strain - PT12123a (referred to as GPSC6wt). PT12123a was isolated from pneumococcal colonization surveillance in a Portuguese day-care center in 2019 (11). It encodes the PhrA1.2-TprA1.2 pair, is genetically tractable, and represents serotype 11A, which is currently expanding in Portugal and Spain (11, 52). As a positive control, we used *S. mitis* C22 where we have recently described PhrA-mediated signaling (Ferreira et al., submitted); as a negative control we used an isogenic deletion mutant of *tprA1.2* (GPSC6Δ*tprA1.2*).

Previous work has shown that synthetic PhrA peptides, corresponding to the C-terminal 10 residues (C-10), activate TprA signaling (28, 40). Thus, to evaluate TprA activity, we measured both autoinduction and expression of the downstream *sce* locus using C-10 PhrA peptides. Neither C-10 PhrA1.1 nor C-10 PhrA1.2 induced *sce* expression in GPSC6wt (**Fig. 2A**), indicating that PhrA signaling is inactive in this background. To determine whether this phenotype reflected a defective peptide or receptor, we reconstructed a full-length *tprA1.2* allele by replacing its premature stop codons with glutamine residues, as found in the majority of TprA alleles (Ferreira et al., submitted). Whole-genome sequencing confirmed the correct replacement of the *tprA1.2* stop codons in the engineered strain **(Table S7).** However, even this full-length variant failed to respond to synthetic PhrA1.2, suggesting that PhrA1.2 does not activate its cognate regulator in GPSC6.

**Figure 2.**
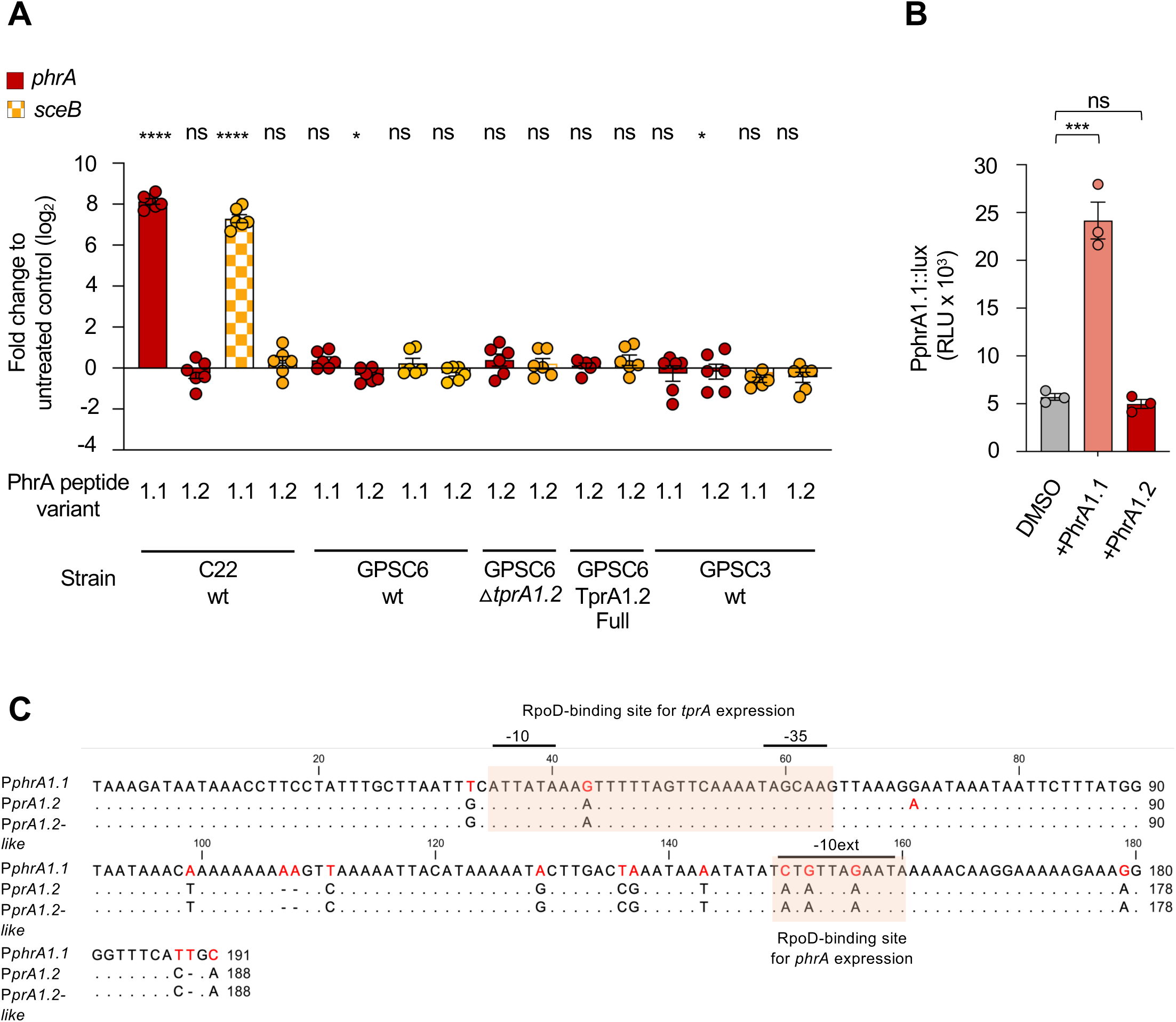
PhrA1.2 is a non-signaling peptide, and *phrA* promoter variation can influence TprA/PhrA-dependent gene regulation in *S. pneumoniae*. A. PhrA-dependent signaling was evaluated by qRT-PCR in strains C22wt, GPSC6wt, GPSC6△*tprA1.2,* GPSC6 TprA1.2-Full and GPSC3wt, in CDM supplemented with galactose. At OD_600nm_ of 0.5, synthetic peptides were added (1mM PhrA1.1 C10 or PhrA1.2 C10), and incubation continued for 2 hours. Gene functions are color-coded consistent with locus diagrams: red, *phrA*; checkered yellow, *sceB* (immunity protein). All assays were performed in triplicates, with two biological replicates per assay. Data represent mean ± SEM. P-values were calculated by paired Student’s t-test. **B.** To assess TprA/PhrA signaling activity, bioluminescence from a *PphrA:luxABCDE* reporter fusion was measured in *S. pneumoniae* D39Δ*phrA* strain. For that, cultures were grown in liquid media and stimulated with 1 mM synthetic PhrA1.1 C10, PhrA1.2 C10, or left untreated (DMSO control). Bioluminescence was monitored over 6 hours. Conditions are color-coded: grey (DMSO), pink (PhrA1.1 C10), red (PhrA1.2 C10). P-values were calculated by two-way ANOVA. (**C**) Alignment of the intergenic regions between *tprA-phrA* for PhrA promoters in strain D39 (with *PphrA1.1*), GPSC6wt (with *PphrA1.2*), and GPSC3 (with *PphrA1.2-like*). Putative key regulatory elements are annotated. \**P* <0.05; \*\*\**P* <0.001; **** P <0.0001; ns, not significant.

Database searches revealed no naturally occurring isolates harboring a full-length TprA1.2, but we identified a highly similar variant, designated TprA1.2-like, differing by two amino acids (S2N and A286T), found in the GPSC3 lineage. Expression analysis of the genes downstream of TprA in a representative strain of this background showed that neither C-10 PhrA1.1 nor C-10 PhrA1.2 activated signaling through this receptor (**Fig 2A**).

Since PhrA1.1 is known to activate the TprA1.1 system in the model strain D39 (28, 40), we next examined whether C-10 PhrA1.2 could relieve TprA-mediated repression using a D39 reporter strain. This construct carries a Δ*phrA* allele in which the *phrA* promoter (PphrA1.1) drives *luxABCDE* expression, requiring exogenous peptide for activation. As expected, C-10 PhrA1.1 induced a >4-fold increase in PphrA1.1 activity; C-10 PhrA1.2, however, had no effect (**Fig. 2B**).

Together, these results demonstrate that PhrA1.2 fails to activate signaling through any tested receptor variant, including the native truncated TprA1.2, an engineered full-length TprA1.2, the closely related TprA1.2-like receptor from GPSC3, and the functionally validated TprA1.1 allele.

As PhrA1.2 failed to activate signaling through any tested receptor variant, we next examined possible molecular bases for its inactivity. We considered three, not mutually exclusive, molecular possibilities for the absence of PhrA1.2 activity. The first is that it does not bind the cognate regulator, the second is that it binds the regulator but that it co-occurs with a promoter that is no longer responsive, and the third is that it is not imported by Ami importer. The latter is unlikely given that the Ami importer is highly promiscuous and imports a large array of peptides with diverse sequence (26). Bioinformatic analysis of the *tprA/phrA* promoter regions (*PphrA*) in GPSC6 (*PphrA1.2*) and GPSC3 (*PphrA1.2-like*) revealed substantial divergence from the corresponding region in strain D39 (*PphrA1.1*). Relative to D39, the GPSC6 and GPSC3 promoters differ by 24 and 23 nucleotides, respectively, and by only a single nucleotide from each other (**Fig. 2C**). These differences raise the possibility that promoter sequence variation contributes to the lack of TprA1.2 activity observed in GPSC6. Because all tested genetic configurations of this system were inactive in GPSC6, we did not experimentally distinguish among potential molecular mechanisms.

### The TprA1.2 provides a fitness advantage *in vitro* and *in vivo* and plays a role in oxidative stress resistance

TprA1.2 does not respond to PhrA peptides yet, it encodes a complete DNA-binding domain. This configuration raises the question as to whether it remains functionally active but peptide independent. To test for this possibility, we generated a *tprA1.2* deletion mutant in GPSC6 strain and compared its growth and global gene expression to that of the parental strain.

We compared the growth profiles of WT and Δ*tprA1.2* strains. Deletion of *tprA1.2* resulted in a growth delay, characterized by an extended lag phase (**Fig. 3A**). This phenotype suggests that TprA1.2 is functionally active in the GPSC6 background. The full-length TprA1.2 (TprA1.2 Full) mimicked the wild-type, suggesting that rescuing the predicted peptide-binding domain did not alter its function.

**Figure 3.**
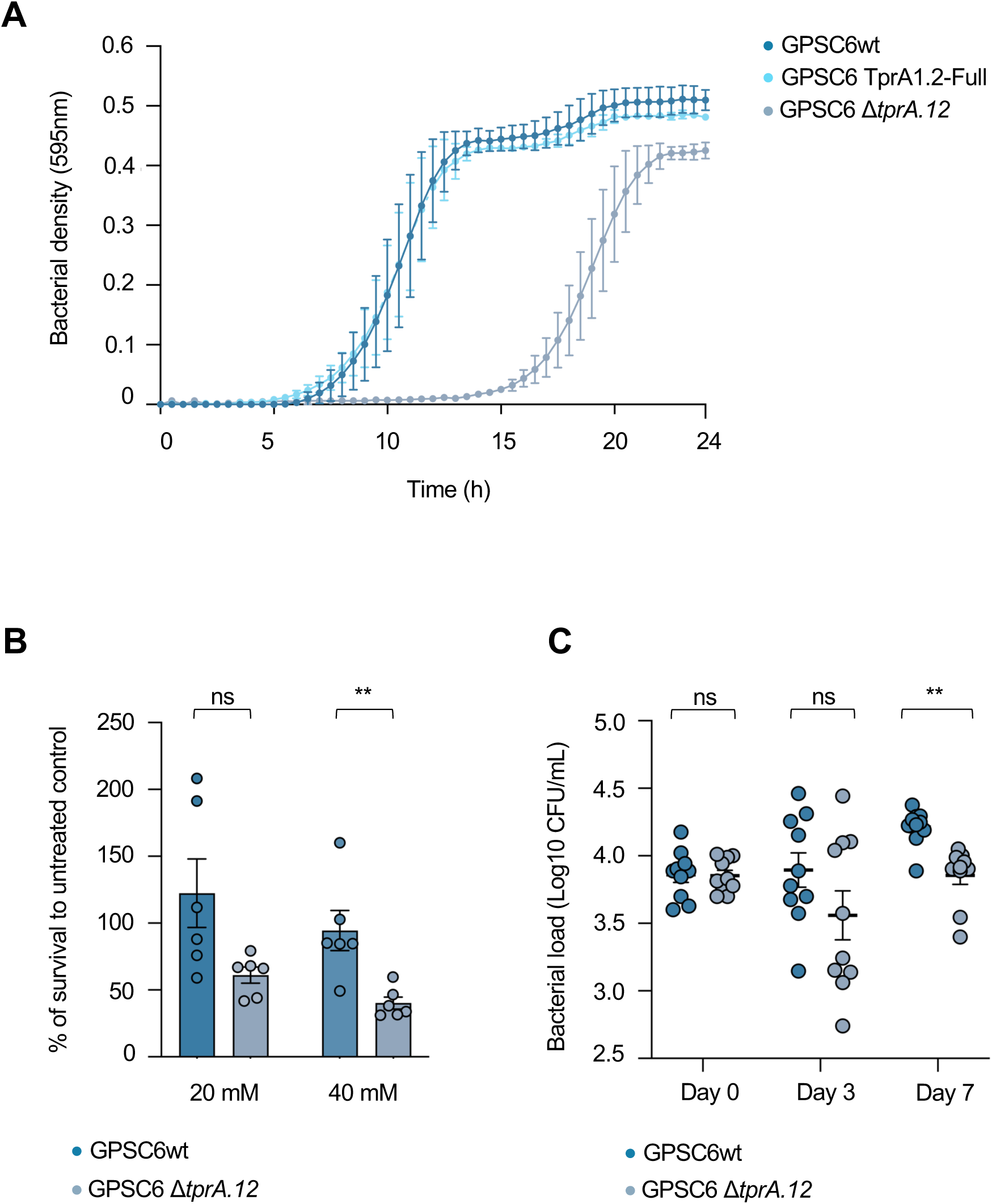
The TprA1.2 allelic variant in GPSC6 lineage enhances bacterial growth, regulates oxidative-stress response, and increases colonization fitness. **A.** Growth of GPSC6wt, GPSC6 TprA-Full, and GPSC6 Δ*tprA1.2* strains was monitored in CDM galactose medium at 37 °C for 24 hours. Assays were performed in triplicate. Data represent the mean ± SEM. **B.** The impact of TprA1.2 on oxidative stress response was assessed by exposing GPSC6wt, and GPSC6Δ*tprA1.2* strains to hydrogen peroxide. Cultures were grown in CDM galactose to an OD_600nm_ of 0.3, treated with 20 mM or 40 mM H_2_O_2_ or left untreated. After 30 minutes, cells were serially diluted and plated on blood agar to quantify survival. Strains are color-coded: dark blue (GPSC6wt) and grey (GPSC6Δ*tprA1.2*). The percentage survival represent the mean ± SEM. P-values were calculated by paired Student’s t-test. (**C**) The contribution of TprA1.2 to nasopharyngeal colonization was evaluated using a murine infection model. Mice (n=10 per strain) were intranasally inoculated with 5×10⁵ CFU of GPSC6wt and GPSC6Δ*tprA1.2*. Bacterial burdens were measured from nasopharyngeal washes at 3- and 7-days post-inoculation. Individual data points and group means are shown. Strains are color-coded: dark blue (GPSC6wt) and grey (GPSC6Δ*tprA1.2*). P-values were calculated by paired Student’s t-test. \**P* <0.05; ns, not significant. \**P* <0.05; \*\**P* <0.001; ns, not significant.

To further assess the role of TprA1.2, we performed RNA-seq to compare global gene expression in the WT and Δ*tprA1.2* strains. Analysis of the TprA1.2 regulon, using a cutoff of absolute log_2_ fold change >1, FDR < 0.05 and p-values <0.001, revealed 5 upregulated and 2 downregulated genes (**Table 1**). Notably, these genes are associated with oxidative stress. They include the *sufCDS* operon, involved in Fe-S cluster biogenesis (53), the predicted ergothioneine transporter *egtV* (54), and *pn-aqpA*, encoding a peroxiporin linked to hydrogen peroxide efflux (55). Collectively, these genes point to a role for TprA1.2 in coordinating the oxidative stress response, suggesting that this regulator contributes to cellular defense against oxidative damage.

**TABLE 1.** Differentially expressed genes in strain GPSC6Δ*tpra1.2* compared to WT.

| Locus tag<br>GPSC6 | Locus<br>tag R6 | Log<sub>2</sub>FC | FDR | p-value | Gene/<br>Function | Genetic control<br>by TprA1.2 |
| --- | --- | --- | --- | --- | --- | --- |
| AIDCGDPD_01078 | - | -8.26 | 4.4E-69 | 1.97e-72 | *tprA1.2* | (deletion made) |
| AIDCGDPD_00828 | spr0772 | -1.21 | 1.9E-03 | 2.64E-06 | *sufD* | Activation |
| AIDCGDPD_00827 | spr0773 | -1.19 | 0.04 | 2.57E-04 | *sufS* | Activation |
| AIDCGDPD_00829 | spr0771 | -1.02 | 0.01 | 3.33E-05 | *sufC* | Activation |
| AIDCGDPD_00692 | spr1604 | -1.05 | 0.01 | 2.57E-05 | *aqpZ* | Activation |
| AIDCGDPD_00907 | spr1678 | -1.01 | 0.04 | 2.56E-04 | *egtV* | Activation |
| AIDCGDPD_01586 | spr0089 | 1.40 | 0.01 | 4.06E-05 | *ptvR* | Repressed |
| AIDCGDPD_00816 | spr0782 | 1.49 | 0.02 | 8.64E-05 | Acetyl-<br>transferase | Repressed |

Cut-off thresholds for gene expression significance: FDR < 0.05, absolute log<sub>2</sub>fold change > 1, and P-value < 0.001.
FDR - false discovery rate.

The set of TprA1.2-controlled genes also included the transcriptional repressor *ptvR* and an additional gene of unknown function (**Table 1**). The PtvR has been shown to repress the downstream *ptvABC* operon, which is associated with vancomycin resistance (56). However, in our conditions, the *ptvABC* expression was unchanged, and we therefore did not investigate potential effects on vancomycin susceptibility.

To determine whether TprA1.2-regulated genes interact at a network level, the differently expressed genes were mapped to the *S. pneumoniae* R6 STRING interactome (**Fig. S1)**. This analysis showed that the Fe-S biogenesis pathway (*sufCDS*) is the only clearly affected multi-gene unit. The remaining TprA1.2-regulated genes did not form a single shared interaction cluster, but their known functions are consistent with adaptation to oxidative stress (e.g. peroxide handling and redox-protective solute transport). Finally, although *ptvR* lies within a regulatory pathway of interest in STRING, its downstream *ptvABC* targets were unchanged under the conditions tested.

Given that TprA1.2 regulates genes linked to oxidative stress tolerance, we performed hydrogen peroxide susceptibility assays in the WT and Δ*tprA1.2* strains (**Fig. 3B**). After 30 minutes of H_2_O_2_ exposure, the WT strain exhibited significantly higher survival than the deletion mutant, supporting a protective role for TprA1.2 under oxidative stress conditions.

To assess the role of TprA1.2 *in vivo*, we employed the murine carriage model of pneumococcal colonization. GPSC6 successfully colonized the murine nasopharynx for at least seven days, allowing us to test how deletion of *tprA1.2* influences colonization density. Bacterial burdens were measured at 3- and 7-days post-inoculation (**Fig. 3C**). The Δ*tprA1.2* strain showed comparable density to the WT at day 0, but significantly lower levels at days 3 and 7. These *in vivo* findings mirror our *in vitro* observations and suggest that TprA1.2 contributes to colonization of GPSC6, potentially through enhanced tolerance to oxidative stress. Together, these data establish TprA1.2 as a regulator that enhances colonization fitness of GPSC6 *in vivo*.

## DISCUSSION

Our work identified and characterized a novel allelic pair of the Tpr/Phr system, TprA1.2/ PhrA1.2, which is strongly associated with the ST156 clone of the highly successful GPSC6 lineage. We find that the PhrA1.2 does not activate signaling. In contrast, the TprA1.2 is a functional regulator, which activates genes linked to oxidative stress responses in a Phr-independent manner. This regulation translates into phenotypic effects, specifically enhanced hydrogen peroxide tolerance and increased colonization capacity in a murine carriage model. These findings illustrate how modifications in cell-cell communication systems can rewire regulatory networks and raise the question of whether such changes have contributed to the epidemiological success of GPSC6 (ST156).

Our analysis of the PhrA peptide revealed that a single amino acid substitution (G7V; position −4) renders the peptide inactive across multiple pneumococcal genetic backgrounds, highlighting the critical role of this position in TprA recognition. The substitution of glycine with valine likely disrupts peptide–receptor interactions within the TprA binding pocket. We propose that glycine’s small, flexible side chain allows the specific conformation required for activation, whereas the bulkier β-branched valine introduces steric hindrance or prevents the peptide from adopting the active conformation. Although direct binding measurements were not performed, the near-complete loss of induction in the PhrA1.2 variant is consistent with impaired receptor recognition or failure to trigger activation. Similar sensitivity to single-residue substitutions has been described in other RRNPP systems, such as PlcR–PapR, NprR–NprX, and ComR-XIP, where peptide-receptor specificity relies on a few residue-dependent sidechain interactions within the TPR-domain pocket (57, 58).

In addition to the loss of signaling by the peptide, our study points to promoter architecture as a potential key determinant of signaling output. In both GPSC6 and GPSC3 isolates, the presence of full-length TprA and PhrA failed to induce substantial downstream gene expression, consistent with cis-regulatory divergence that can silence a CCC module without requiring protein-coding changes. In line with this, TprA1.2 no longer regulates the *sce* operon, suggesting that SNPs accumulated on the promoter have disrupted TprA-DNA interaction. Notably, a similar loss of responsiveness is observed in other genetic backgrounds, including GPSC3, indicating that decay of the TprA-DNA interaction in the control of *sce* expression may have occurred independently across pneumococcal lineages or been horizontally transferred across lineages.

Despite this loss of inhibition at the *sce* locus, our transcriptomics data demonstrate that TprA1.2 retains regulatory activity at other genomic loci, acting as an activator of genes involved in oxidative stress defense, including Fe–S cluster biogenesis, redox homeostasis, and hydrogen peroxide export (53–55). These functions persist despite the absence of a complete PhrA-binding domain, indicating that transcriptional activation has become uncoupled from peptide-mediated signaling. Similar rewiring events have been documented in other bacterial communication systems (59, 60). Compared to the TprA regulon of the reference pneumococcal strain D39 (28), our data reveal a functional shift: whereas TprA1.1 in D39 regulates antimicrobial peptide resistance and carbohydrate uptake, TprA1.2 in GPSC6 primarily activates stress-response pathways. This supports the idea that partial loss of the pheromone-binding domain drove a repurposing of TprA activity, redirecting the system from roles in competition and nutrient sensing toward oxidative stress adaptation.

Comparative genomics revealed that the PhrA1.2/TprA1.2 module is not unique to GPSC6 (ST156) but is also present in other pneumococcal lineages, such as GPSC39 (ST124). However, while GPSC6 has retained and functionally integrated the system, GPSC39 shows evidence of progressive loss: older genomes retained the locus, but most recent isolates lack TprA1.2/PhrA1.2. This pattern indicates divergent selective trajectories acting on the same module. In GPSC6, the system appears to have been maintained, likely due to its beneficial role in oxidative stress tolerance and colonization, whereas in GPSC39, it may have become dispensable under distinct ecological or epidemiological conditions.

From an evolutionary and epidemiological perspective, these contrasts underscore how cell-cell communication modules can follow lineage-specific evolutionary paths, being maintained, silenced, or lost depending on their adaptive value. In GPSC6, it is tempting to speculate that the functional retention and repurposing of PhrA1.2/TprA1.2 may have contributed to the lineage’s persistence, transmission efficiency, and global dissemination, alongside its well-known multidrug resistance and serotype plasticity.

## ACKNOWLEDGMENTS

We thank Adriano Henriques (ITQB NOVA) and Cristina Silva Pereira (ITQB NOVA) for interesting and productive discussions.

## CONTRIBUTIONS

RSL, CV and LH were responsible for the concept and design of the study. RSL contributed with study collections, reagents and materials. LH and HY contributed with reagents and materials. BF performed all experimental work except for the *in vivo* studies, which were performed by OG. Data analysis and interpretation of results were done by BF, CV, LH and RSL. The manuscript was drafted by BF, CV, LH and RSL and critically revised by all authors. All authors read and approved the final version of the manuscript.

