## Supplementary Figure for "Network rewiring of a pneumococcal communication system promotes stress adaptation in a globally successful lineage"

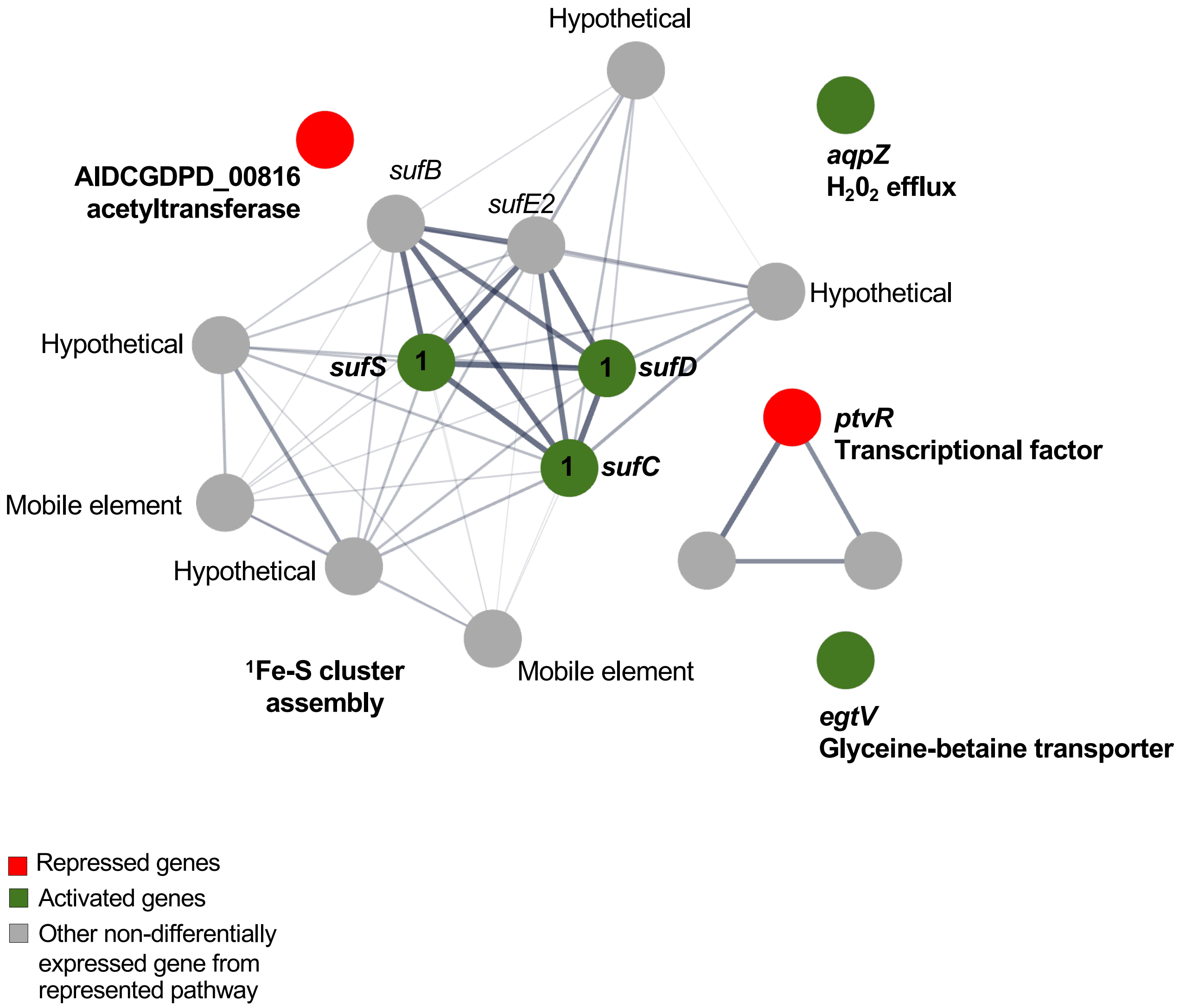


**Figure S1. STRING association network of differentially expressed (DE) genes under TprA1.2 control.** GPSC6 DE genes ($|\log_2 FC| > 1$; FDR < 0.05; $P < 0.001$) were mapped to *S. pneumoniae* R6 orthologues and submitted to STRING. Red nodes: genes repressed by TprA1.2; green nodes: genes activated by TprA1.2. Grey nodes: non-differentially expressed predicted functional partners returned by STRING within the represented modules. Functional modules recovered from the STRING network (e.g. Fe–S cluster assembly, $H_2O_2$ efflux, glycine-betaine transport, transcriptional regulation) are indicated by annotation.
