## Supplementary Tables for "Network rewiring of a pneumococcal communication system promotes stress adaptation in a globally successful lineage"

**Table S1. Strains, plasmids, oligonucleotides and peptides used in the study.**

| Bacterial strain, plasmid, oligonucleotide or peptide | Relevant genotype or sequence | Note | Source |
|---|---|---|---|
| ***Streptococcus pneumoniae* strains** | | | |
| GPSC6wt | Wild-type strain. 11A serotype. PT12123a from Candeias et al. (REF) | | Candeias *et al.*, 2024 |
| GPSC6 Δ*tprA1.2* | Δ*tprA1.2* markerless mutant | | This study |
| GPSC6 TprA1.2-full | *tpra1.2*$^{*216Q,*229Q}$ | | This study |
| GPSC3wt | Wild-type strain. 33F serotype. PT12230 | | |
| D39PphrA:lux | Cassette insertion in *bgaA* non coding region of genome | | Muller-Brown et al., 2024 |
| D39 ΔphrA PphrA:lux | Cassette insertion in *bgaA* non coding region of genome | | Muller-Brown et al., 2024 |
| **Oligonucleotides for the construction of GPSC6 TprA1.2-full** | | | |
| P1F_TprA_Replacement | CAAGTGAATATCTAGGTGTC | Reversion of early stop codons from *tprA1.2* from GPSC6: tpra1.2*216Q,*229Q | This study |
| P1RTprA_Replacement_Fin | TTGCCGCACTGATAGAGTATTCAACGCC | | |
| P2F_Franc_TprA_Replacement_Fin | ATACTCTATCAGTGCGGCAAGCCAAGCA | | |
| P2R_Franc_TprA_Replacement_Fin | CTCCAGCATTCCTGTCTTTGGGCAGATAAAGGC | | |
| P3F_TprA_Replacement_Fin | CAAAGACAGGAATGCTGGAGCCAAATTTC | | |
| P3R_TprA_Replacement | GGGCTCTGCTTCCCTAATG | | |
| P1R__TprA_Replacement_Fout | GATAGAGTATTCAACGCC | | |
| P2F_TprA_Replacement_Fout | AAGGCGTTGAATACTCTATC | | |
| P2R_TprA_Replacement_Fout | TGACTTGTTCTCTGTCTAG | | |
| P3F_TprA_Replacement_Fout | AATGCTGGAGCCAAATTTC | | |
| **Oligonucleotides for the construction of GPSC6 Δ*tprA1.2*** | | | |
| P1F_TprAdeletion | ACGCTCCAAGTAATGCGG | Deletion of tprA1.2 from GPSC6wt | This study |
| P1R_TprAdeletion_Fin | TTGCCGCACTCTCTGCGAGTGTACTCATTAAAG | | |

| | | | |
|---|---|---|---|
| P2F_Franc_TprAdeletion_Fin | ACTCGCAGAGAGTGCGGCAAGCCAAGCA | | |
| P2R_Franc_TprAdeletion_Fin | TAAACTAAACCCTGTCTTTGGGCAGATAAAGGC | | |
| P3F_TprAdeletion_Fin | CAAAGACAGGGTTTAGTTTATCTTGATATTGGAGTG | | |
| P3R_TprAdeletion | CGAACTAGCTTGCTTGTTG | | |
| P1R_TprAdeletion_Fout | TAAACTAAACCTCTGCGAGTGTACTCATTAAAG | | |
| P2F_TprAdeletion_Fout | ACTCGCAGAGGTTTAGTTTATCTTGATATTGGAGTG | | |
| **Oligonucleotides for RT-qPCR** | | | |
| PF_TprA_GPSC6wt | GTTGAGACTCTCCCAACAAACT | Gene *tprA1.2* | This study |
| PR_TprA_GPSC6wt | TCTGCGGAGGGAATGAAATG | | |
| PF_PhrA_GPSC6 | TCCTTACCAACAGTTTATTCAGAAG | Gene *phrA1.2* | This study |
| PR_PhrA_GPSC6 | CCTTCCCAACATCAAGACCA | | |
| PF_IM_StrainC22 | ACTAGCAGAAGAAACGGATAGC | Gene *sceB* | Ferreira *et al.*, submitted |
| PR_IM_StrainC22 | CCGTTCCCAAAGGTCGTATAA | | |
| PF_16S | ACCCGAAGTCGGTGAGGTA | Gene *16S* (reference gene) | Muller-Brown *et al.*, 2024 |
| PR_16S | CCAAATCATCTATCCCACCTT | | |
| **Synthetic PhrA peptides** | | | |
| PhrA1.1 C10 | NH2-SNGLDVGKAD-COOH | | This study |
| PhrA1.2 C10 | NH2-SNGLDVVKAD-COOH | | |
| **Synthetic CSP peptide** | | | |
| CSP1 | NH2-EMRLSKFFRDFILQRKK-COOH | | This study |

**Table S2. Genomic context of PhrA1.2 across 7,548 S. pneumoniae isolates**

<table>
<tr><th>PhrA allele</th><th>TprA allele</th><th>TprA sequence (aa)</th><th>Downstream genetic region</th><th>Isolation years</th><th>GPSC profile</th><th>No. of isolates</th></tr>
<tr><td rowspan="6">PhrA1.2</td><td>TprA1.2</td><td rowspan="2">MSTLAEKFRLKRKELRLSQQTLAEGICEQSQISKIERGHFIPSADLLFKLSQ<br>RLEVPLDYFFNEQIEIKSNLSNFKQLSARLLDDRNYDDLEYIYRIEIERSTF<br>LTLEDRTYLEWIKAIIDFYQYDSKCEAISSLENILLKVSSNTLIYLKALNTL<br>SNFYSLVGREQEYEANYSHLIELYQTKNLDHQEFLFGYIRVRYNYAHYLVSK<br>EKYNEAI*EALETIELCKQR*TSYQLAPLLILVGNAGAKFLDREQVKNYYIE<br>ARELCKIYNNPLMLMKIENYLKELDAV*</td><td rowspan="6">Streptococcin E incomplete locus (sceBC operon)</td><td>1962-2015</td><td>6</td><td>161</td></tr>
<tr><td>TprA1.2</td><td>1952</td><td>39</td><td>3</td></tr>
<tr><td>TprA1.3</td><td>MSTLAEKFRLKRKELRLSQQTLAEGICEQSQISKIERGHFIPSADLLFKLSQ<br>RLEVPLDYFFNEQIEIKSNLSNFKQLSARLLDDRNYDDLEYIYRIEIERSTF<br>LTLEDRTYLEWIKAIIDFYQYDSKCEAISSLENILLKVSSNTLIYLKALNTL<br>SNFYSLVGREQEYEANYSHLIELYQTKNLDHQEFLFGYIRVRYNYAHYLVSK<br>EKYNEAI*EALETIELCKQRQTSYQLAPLLILVGNAGAKFLDREQVKNYYIE<br>ARELCKIYNNPLMLMKIENYLKELDAV*</td><td>2007</td><td>268</td><td>4</td></tr>
<tr><td>TprA1.4</td><td>MSTLAEKFRLKRKELRLSQQTLAEGICERSQISKIERGHFIPSADLLFKLSQ<br>RLEVPLDYFFNEQIEIKSNLSNFKQLSARLLDDRNYDDLEYIYRIEIERSTF<br>LTLEDRTYLEWIKAIIDFYQYDSKCEAISSLENILLKVSSNTLIYLKALNTL<br>SNFYSLVGREQEYEANYSHLIELYQTKNLDHQEFLFGYIRVRYNYAHYLVSK<br>EKYNEAI*EALETIELCKQR*TSYQLAPLLILVGNAGAKFLDREQVKNYYIE<br>ARELCKIYNNPLMLMKIENYLKELDAV*</td><td>2011</td><td>6</td><td>2</td></tr>
<tr><td rowspan="2">-</td><td rowspan="2">-</td><td>2015</td><td>6</td><td>2</td></tr>
<tr><td>NA</td><td>2009</td><td>1</td></tr>
</table>

NA - not assigned

**Table S3. Epidemiological context, within TprA/PhrA framework, of GPSC6, GPSC39 and GPSC268 across 7,548 S. pneumoniae isolates.**

| GPSC profile | ST | PhrA allele | TprA allele | Year(s) isolation | No. of isolates |
|---|---|---|---|---|---|
| 6 | 143 | PhrA1.2 | TprA1.2 | 2009 | 3 |
| | 156 | Not assigned[a] | TprA1.195 | 2013 | 1 |
| | | PhrA1.2 | No tprA | 2009-2015 | 2 |
| | | No phrA | No tprA | 1993-2010 | 3 |
| | | PhrA1.2 | TprA1.2 | 1987-2015 | 51 |
| | 162 | PhrA1.2 | TprA1.124 | 2011 | 2 |
| | | PhrA1.2 | TprA1.2 | 1991-2014 | 76 |
| | 312 | PhrA1.2 | TprA1.2 | 1962 | 1 |
| | 1206 | PhrA1.2 | TprA1.2 | 1999-2000 | 2 |
| | 1269 | No phrA | TprA1.16 | 2009-2010 | 6 |
| | | PhrA1.2 | TprA1.2 | 1998-2007 | 10 |
| | 1925 | No phrA | TprA1.5 | 2004-2007 | 4 |
| | 2306 | PhrA1.2 | TprA1.2 | 2010-2011 | 4 |
| | 3057 | PhrA1.2 | TprA1.2 | 1998 | 3 |
| | 3275 | PhrA1.2 | TprA1.2 | 2007 | 1 |
| | 4026 | PhrA1.2 | TprA1.2 | 2006 | 1 |
| | 4464 | PhrA1.2 | TprA1.2 | 2005 | 1 |
| | 8138 | PhrA1.2 | TprA1.2 | 2007 | 1 |
| | 8398 | PhrA1.2 | TprA1.2 | - | 4 |
| | 10449 | PhrA1.2 | TprA1.2 | 2011 | 1 |
| | NA | PhrA1.2 | TprA1.2 | 1998-2003 | 2 |
| 39 | 124 | PhrA1.3 | TprA1.175 | 2009 | 1 |
| | | PhrA1.2 | TprA1.2 | 1952 | 3 |
| | | PhrA1.3 | TprA1.5 | 1980-2012 | 14 |
| | | No phrA | TprA1.17 | 1978-2014 | 69 |
| | 134 | PhrA1.1 | TprA1.11 | 1961 | 1 |
| | 656 | No phrA | TprA1.17 | 1998 | 1 |
| | 2194 | No phrA | TprA1.17 | 1999 | 2 |
| | 3049 | No phrA | TprA1.17 | 1998 | 5 |
| | 5979 | PhrA1.1 | TprA1.11 | 1952 | 1 |
| | 7198 | No phrA | TprA1.17 | 1992 | 1 |
| | 7201 | PhrA1.1 | TprA1.19 | 1957 | 1 |
| | 7253 | No phrA | TprA1.17 | 1989 | 1 |
| | 9683 | No phrA | TprA1.17 | 1989 | 1 |
| | 13127 | No phrA | TprA1.17 | 2012 | 1 |
| 268 | NA | PhrA1.2 | TprA1.85 | 2007 | 4 |

GPS - global pneumococcal sequence cluster

ST - sequence type

[a] No PhrA variant was atributted given it was only found in this genome

NA - not assigned

**Table S4. MLST profiles and genetic relatedness among GPSC6 isolates with PhrA1.2 and TprA1.2 from Golden dataset**

| ST | *aroE* | *gdh* | *gki* | *recP* | *spi* | *xpt* | *ddl* | Locus variants | Number of. isolates |
|---|---|---|---|---|---|---|---|---|---|
| 156 | 7 | 11 | 10 | 1 | 6 | 8 | 1 | Reference | 51 |
| 143 | 7 | 5 | 10 | 18 | 6 | 8 | 1 | DLV | 3 |
| 162 | 7 | 11 | 10 | 1 | 6 | 8 | 14 | SLV | 76 |
| 312 | 7 | 11 | 10 | 1 | 6 | 1 | 14 | DLV | 1 |
| 1206 | 7 | 11 | 41 | 1 | 6 | 8 | 14 | DLV | 2 |
| 1269 | 7 | 11 | 10 | 1 | 6 | 76 | 14 | DLV | 10 |
| 2306 | 7 | 11 | 10 | 1 | 6 | 8 | 119 | SLV | 4 |
| 3057 | 7 | 11 | 41 | 1 | 6 | 161 | 14 | TLV | 3 |
| 3275 | 7 | 183 | 10 | 1 | 6 | 8 | 14 | DLV | 1 |
| 4026 | 7 | 11 | 10 | 1 | 6 | 8 | 316 | SLV | 1 |
| 4464 | 7 | 11 | 10 | 1 | 6 | 198 | 1 | SLV | 1 |
| 8138 | 7 | 11 | 10 | 16 | 6 | 493 | 14 | TLV | 1 |
| 8398 | 7 | 11 | 10 | 1 | 8 | 8 | 1 | SLV | 4 |
| 10449 | 7 | 11 | 10 | 1 | 451 | 8 | 1 | DLV | 1 |

SLV - single-locus variant; DLV - double-locus variant; TLV - triple-locus variant
In green is highlighted allele difference towards the reference (ST156)

**Table S5. Genomic context of PhrA1.2 across 873 *S. pneumoniae* isolates from Portuguesse carriage study**

<table>
<tr><th>PhrA allele</th><th>TprA allele</th><th>TprA sequence (aa)</th><th>Downstream genetic region</th><th>Isolation years</th><th>GPSC profile</th><th>No. of isolates</th></tr>
<tr><td rowspan="2">PhrA 1.2</td><td rowspan="2">TprA 1.2</td><td rowspan="2">MSTLAEKFRLKRKELRLSQQTLAEGICEQSQISKIERGHFIPSADLLFKLSQRLEVPLDYFFNEQIEIKSNLSNFKQLSARLLDDRNYDDLEYIYRIEIERSTFLTLEDRTYLEWIKAIIDFYQYDSKCEAISSLENILLKVSSNTLIYLKALNTLSNFYSLVGREQEYEANYSHLIELYQTKNLDHQEFLFGYIRVRYNYAHYLVSKEKYNEAI*EALETIELCKQR*TSYQLAPLLILVGNAGAKFLDREQVKNYYIEARELCKIYNNPLMLMKIENYLKELDAV*</td><td rowspan="2">Streptococcin E incomplete locus (sceBC operon)</td><td rowspan="2">2018-2020</td><td>6</td><td>106</td></tr>
<tr><td>NA</td><td>1</td></tr>
</table>

NA - not assigned

**Table S6. MLST profiles and genetic relatedness among GPSC6 isolates with PhrA1.2 and TprA1.2 from Portuguese dataset**

| ST | *aroE* | *gdh* | *gki* | *recP* | *spi* | *xpt* | *ddl* | Locus variants | Number of. isolates |
|---|---|---|---|---|---|---|---|---|---|
| 156 | 7 | 11 | 10 | 1 | 6 | 8 | 1 | Reference | 5 |
| 162 | 7 | 11 | 10 | 1 | 6 | 8 | 14 | SLV | 49 |
| 166 | 7 | 11 | 10 | 1 | 6 | 1 | 1 | SLV | 1 |
| 838 | 7 | 11 | 10 | 1 | 6 | 8 | 90 | SLV | 2 |
| 6521 | 8 | 11 | 10 | 1 | 6 | 8 | 90 | DLV | 39 |
| 14368 | 7 | 11 | 10 | 1 | 6 | 1 | 930 | DLV | 12 |

SLV - single-locus variant; DLV - double-locus variant; TLV - triple-locus variant

In green is highlighted allele difference towards the reference (ST156)

**Table S7. Single nucleotide polymorphisms detected in the comparasion of GPSC6 TprA-Full and GPSC6wt**

| Annotation in reference pneumococcal strain[a] | Gene name | Genome region | Nucleotide change[b] | Amino acid change | Frequency (%) | Average quality |
|---|---|---|---|---|---|---|
| SPD_1649 | *piuB* | Contig 6 | C36610A | Glu208* | 100 | 33.7 |
| SPD_1745 | *tprA1.2* | Contig 8 | T51212C | ***216Gln** | 100 | 33.8 |
| SPD_1745 | *tprA1.2* | Contig 8 | T51251C | ***229Gln** | 100 | 33.8 |
| SPD_0510 | *metE* | Contig 13 | G11983T | **Ala95Ser** | 100 | 31.6 |

[a]D39 (SPD_);

[b]Nucleotide changes with minimum frequency of 95% and average base quality ≥20;

[c]Amino acid changes in bold are putatively deleterious according to SIFT web server, or by search of lost domains in InterProScan and NCBI
